# Sequence structure organizes items in varied latent states of working memory neural network

**DOI:** 10.1101/2020.06.20.162438

**Authors:** Qiaoli Huang, Huihui Zhang, Huan Luo

## Abstract

In memory experiences, events do not exist independently but are linked with each other via structure-based organization. Structure context largely influences memory behavior, but how it is implemented in the brain remains unknown. Here, we combined magnetoencephalogram (MEG) recordings, computational modeling, and impulse-response approaches to probe the latent states when subjects held a list of items in working memory (WM). We demonstrate that sequence context reorganizes WM items into distinct latent states, i.e., being reactivated at different latencies during WM retention, and the reactivation profiles further correlate with recency behavior. In contrast, memorizing the same list of items without sequence task requirements disrupts the recency effect and elicits comparable neural reactivations. Computational modeling further reveals a dominant function of sequence context, instead of passive memory decaying, in characterizing recency effect. Taken together, sequence structure context shapes the way WM items are stored in the human brain and essentially influences memory behavior.

## Introduction

Working memory (WM), more than just passively maintaining inputs, is also an active process that modifies and even reorganizes information representations to guide future behavior (Baddeley, 2003). For instance, retro-cues during retention could enhance WM performance (Griffin & Nobre, 2003; Landman et al., 2003; LaRocque et al., 2014) and modulate neural responses in different brain regions (Christophel et al., 2018; Yu et al., 2020), suggesting that attention could flexibly modulate information that has already been maintained in WM (Myers et al., 2017). In addition to top-down attention, contexts and structure in which the to-be-memorized items are embedded also influence memory performance (Brady et al., 2011; Brady & Tenenbaum, 2013; DuBrow & Davachi, 2013; Gershman et al., 2013; Jiang et al., 2000; Oberauer & Lin, 2017). A typical example is the serial position effect for sequence memory, i.e., the recently presented item shows better memory performance compared to early one (Baddeley, 2012; Burgess & Hitch, 1999; Gorgoraptis et al., 2011; Huang et al., 2018; Jones & Oberauer, 2013). However, it remains largely unknown how structure context modulates or reorganizes the way multiple WM items are represented and maintained in the human brain. Here we particularly focused on sequence structure, an essential one that mediates many cognitive functions, e.g., sequence memory, speech, movement control, etc (Davachi & DuBrow, 2015; Giraud & Poeppel, 2012; Polyn et al., 2009).

Previous neural recordings demonstrate that a sequence of items would elicit serial and temporally compressed reactivations, which might reflect a memory consolidation process that reorganizes and stores the incoming inputs (Bahramisharif et al., 2018; D. J. Foster & Wilson, 2006; Huang et al., 2018; Kurth-Nelson et al., 2016; Y. Liu et al., 2019; Siegel et al., 2009; Skaggs & McNaughton, 1996). Interestingly, recent modeling and empirical studies propose that, instead of maintenance via sustained or serial reactivations (Curtis & D’Esposito, 2003; Goldman-Rakic, 1995; Vogel & Machizawa, 2004), information could also be stored in synaptic weights of the network and thus does not rely on sustained responses, i.e., activity-silent view (Miller et al., 2018; Mongillo et al., 2008; Rose et al., 2016; Sprague et al., 2016; Stokes, 2015; Trübutschek et al., 2017; Wolff et al., 2017). In other words, multiple items could be maintained in latent or hidden states of the WM neural network.

How to access the WM information retained in the ‘activity-silent’ network? An impulse-response approach, by presenting a PING stimulus to transiently perturb the WM system, could efficiently reactivate WM representations (Wolff et al., 2017). Moreover, attended but not unattended item is successfully reactivated, implying that top-down attention modulates the latent states WM items reside in (Wolff et al., 2017, 2020). Here we used this method to assess whether a list of WM items of equal task relevance would be reorganized by imposed sequence structure in varied latent states and in turn show distinct reactivation profiles. The hypothesis is also motivated by our previous findings demonstrating backward reactivations for sequence memory, which implies a more excitable state for recent vs. early items (Huang et al., 2018).

We recorded magnetoencephalography (MEG) signals when human subjects (N = 24) retained a sequence of orientations and their ordinal positions. A time-resolved multivariate inverted encoding model (IEM) (Brouwer & Heeger, 2009, 2011; Sprague et al., 2014) was used to reconstruct the neural representations of each orientation (i.e., 1^st^ item, 2^nd^ item) over time. Importantly, a nonspecific PING stimulus (i.e, impulse) was presented during retention, aiming to transiently perturb the WM network so that the latent states of the stored items could be accessed. Our results demonstrate a backward reactivation profile such that the impulse triggers the neural representation of the 2^nd^ item first, followed by that of the 1^st^ item, thus supporting their different latent states. Moreover, the neural reactivation pattern well predicts the recency effect in behavior. In contrast, in another MEG experiment when subjects (N =24, new subjects) retained the same sequence without needing to memorize the ordinal structure, the two orientations show similar reactivation profiles. Finally, computational modelling demonstrates that the sequence contexts, instead of passive memory decay, largely characterizes the recency behavior. Our findings constitute converging evidence supporting a central function of sequence structure in WM via reorganizing multiple items in the brain (i.e., in varied latent states) and influencing memory behavior. Generally speaking, our findings provide new perspectives for the neural mechanism underlying multi-item information storage in the WM system.

## Results

### Experimental procedure and recency behavior (Experiment 1)

Twenty-four subjects participated in Experiment 1 and their brain activities were recorded using a 306-channel magnetoencephalography (MEG) system (Elekta Neuromag system, Helsinki, Finland). As shown in Figure 1A, each trial consists of three periods – encoding, maintaining, and recalling. During the encoding period, participants viewed two serially presented grating stimuli and were instructed to memorize the two orientations as well as their order (1^st^ and 2^nd^ orientations). After a 2 sec maintaining period, a retrospective cue (retro-cue) appeared to instruct subjects which item (1^st^ or 2^nd^) would be tested. Next, a probe grating was presented and participants indicated whether the orientation of the probe was rotated clockwise or anticlockwise relative to that of the cued grating. Note that since the retro-cue appeared only during the recalling period, subjects would need to hold the two WM orientations simultaneously in WM throughout the retention interval. Critically, during the maintaining period, a high-luminance PING stimulus that does not contain any orientation information was presented, aiming to transiently perturb the WM network so that the stored information and its associated latent states could be assessed.

**Figure 1:**
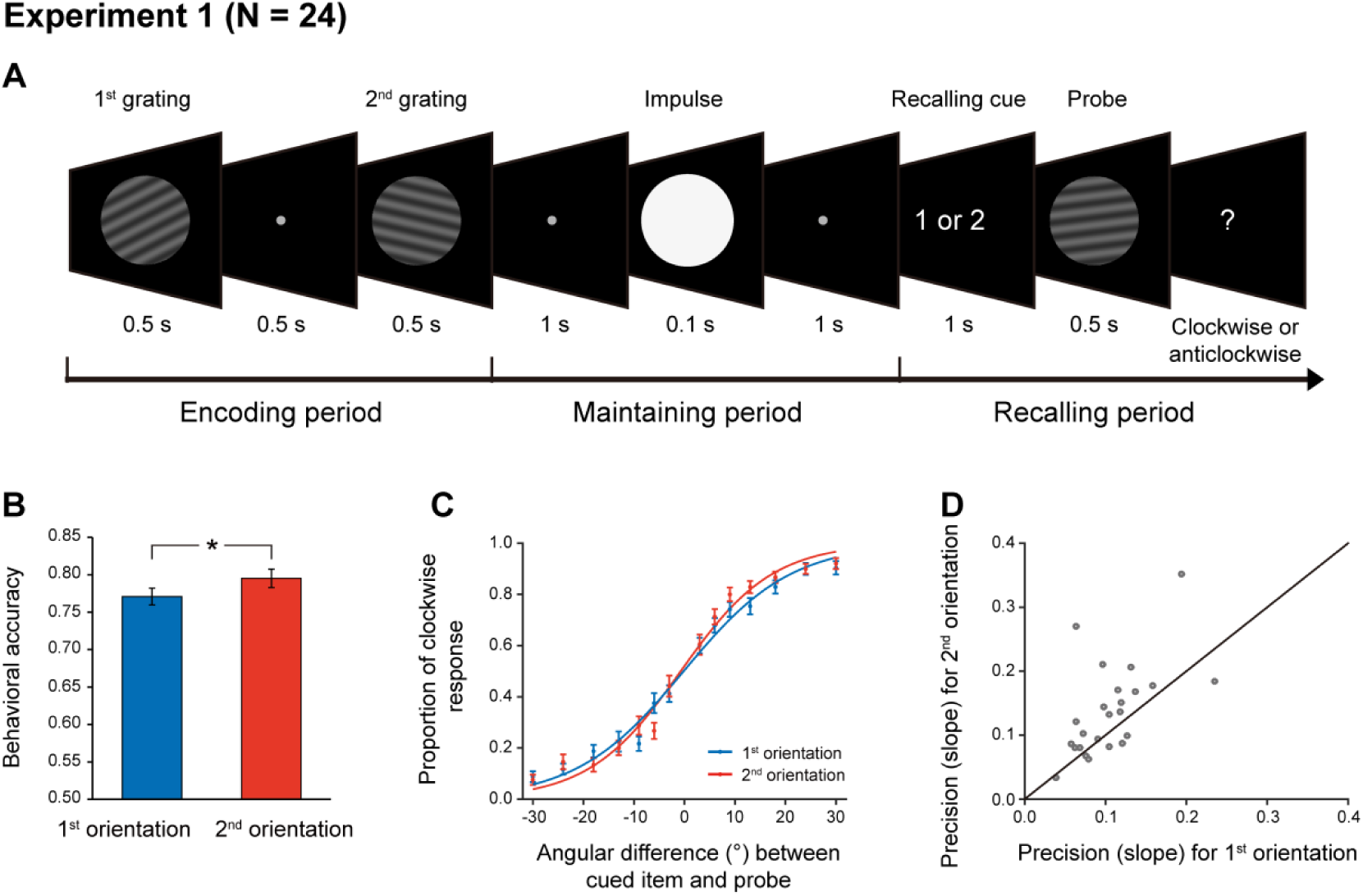
Experimental paradigm and recency effect (Experiment 1). (A) Experiment 1 paradigm (N = 24). Subjects were first sequentially presented with two grating stimuli (1^st^ and 2^nd^ gratings) and needed to memorize the orientations of the two gratings as well as their temporal order (1^st^ or 2^nd^ orientation). During the maintaining period, a high luminance disc that does not contain any orientation information was presented as a PING stimulus to disturb the WM neural network. During the recalling period, a retro-cue first appeared to instruct subjects which item (1^st^ or 2^nd^) would be tested. Next, a probe grating was presented and participants indicated whether the orientation of the probe was rotated clockwise or anticlockwise relative to that of the cued grating. (B) Grand average (mean ± SEM) behavioral accuracy for the 1^st^ (blue) and 2^nd^ (red) orientations. (C) Grand average (mean ± SEM) psychometric functions for the 1^st^ (blue) and 2^nd^ (red) items as a function of the angular difference between the probe and cued orientation. Note the steeper slope for the 2^nd^ vs. 1^st^ orientation, i.e., recency effect. (D) Scatter plot for the slope of the psychometric function for the 1^st^ (x axis) and 2^nd^ orientations (y axis). (*: p < 0.05).

As shown in Figure 1B, the memory behavioral performance exhibited the typical recency effect, i.e., 2^nd^ > 1^st^ item (N = 24, 1^st^ item: mean = 0.77, s.e. = 0.012; 2^nd^ item: mean = 0.79, s.e = 0.013; paired t-test, df = 23, t = 2.18, p = 0.039, Cohen’s d = 0.45), consistent with previous findings (e.g., Baddeley, 2012; Burgess & Hitch, 1999; Gorgoraptis et al., 2011; Huang et al., 2018; Jones & Oberauer, 2013). When plotting the psychometric function of the proportion of clockwise response as a function of angular difference between the probe and the cued WM grating, the 2^nd^ item showed steeper slope than the 1^st^ item (Figure 1C; 1^st^ item: mean = 0.11, s.e. = 0.009; 2^nd^ item: mean = 0.14, s.e = 0.015; paired t-test, df = 23, t = 2.64, p = 0.015, Cohen’s d = 0.54). This pattern could be reliably observed for individual subjects (Figure 1D), consistently supporting the recency effect.

### Time-resolved neural representations of orientation features (Experiment 1)

We used a time-resolved inverted encoding model (IEM) (Brouwer & Heeger, 2009, 2011; Sprague et al., 2014) to reconstruct the neural representations of orientation features at each time point throughout the experimental trial. We first verified this method by applying it to the encoding period when the to-be-memorized grating stimuli were presented physically. Specifically, the orientation decoding performance was characterized by reconstructed channel response as a function of angular difference between an orientation-of-interest and other orientations (see details in Methods). If MEG signals do carry information about specific orientation, the reconstructed channel response would reveal a peak at center and gradually decrease on both sides. Figure 2AB plot reconstructed channel responses for the 1^st^ and 2^nd^ WM orientations, respectively, as a function of time throughout the encoding phase. It is clear that right after the presentation of the 1^st^ grating, the reconstructed channel response for the 1^st^ orientation showed central peak (Figure 2A), whereas that for the 2^nd^ orientation displayed information representation only after the presentation of the 2^nd^ grating (Figure 2B). To further quantify the time-resolved decoding performance, the slope of the reconstructed channel response was estimated at each time point in each trial, for the 1^st^ and 2^nd^ orientations, respectively (see details in Methods). As shown in Figure 2C, the 1^st^ orientation (blue) showed information representation right after the 1^st^ grating (0.05 – 0.39 s, corrected cluster p = 0.002; 0.5 – 0.88 s, corrected cluster p < 0.001; 1.13 – 1.4 s, corrected cluster p = 0.002), and neural representation of the 2^nd^ orientation (red) emerged after the 2^nd^ grating (1.06 – 1.5 s, corrected cluster p < 0.001; 1.57 – 1.86 s, corrected cluster p = 0.008). Therefore, orientation information could be successfully decoded from MEG signals in a time-resolved manner. Moreover, the decoding performance for both the 1^st^ and 2^nd^ orientations gradually decayed to baseline, around 0.5 s after the offset of the 2^nd^ item, suggesting that the WM network now entered the activity-silent WM states (Rose et al., 2016; Stokes, 2015; Wolff et al., 2017). It is notable that nonsignificant decoding results do not exclude sustained firing at neuronal level (Miller et al., 2018), given the limited sensitivity of MEG and EEG signals.

**Figure 2:**
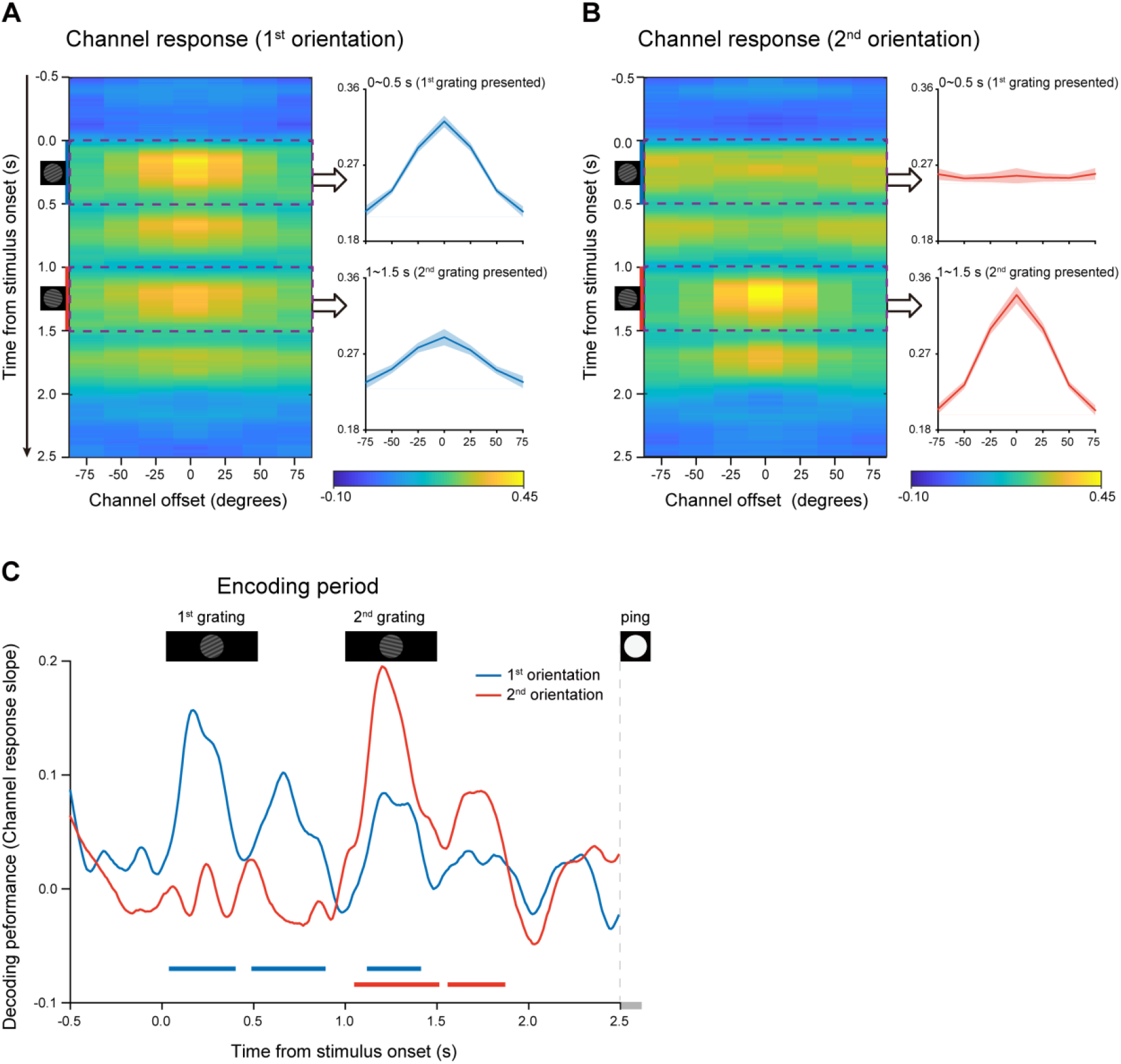
Time-resolved orientation representations during encoding period (Experiment 1). An IEM was used to reconstruct the neural representation of orientation features, characterized as a population reconstructed channel response as a function of channel offset (x-axis) at each time bin (y-axis). Successful encoding of orientation features would show a peak at center around 0 and gradually decrease on both sides, whereas lack of orientation information would have a flat channel response. The slope of the channel response was further calculated (see methods for details) to index information representation. (A) Left: Grand average time-resolved channel response for the 1^st^ orientation (orientation of the 1^st^ WM grating) throughout the encoding period during which the 1^st^ and 2^nd^ gratings (small inset figures on the left) were presented sequentially. Right: grand average (mean ± SEM) channel response for the 1^st^ orientation averaged over the 1^st^ grating presentation period (0 – 0.5 s, upper) and the 2^nd^ grating presentation (1 – 1.5 s, lower). (B) Left: Grand average time-resolved channel response for the 2^nd^ orientation (orientation of the 2^nd^ WM grating) during the encoding period. Right: grand average (mean ± SEM) channel response for the 2^nd^ orientation averaged over the 1^st^ (0 – 0.5 s, upper) and 2^nd^ (1 – 1.5 s, lower) grating presentation period. (C) Grand average time courses of the channel response slopes for the 1^st^ (blue) and 2^nd^ (red) orientations during the encoding period. Horizontal lines below indicate significant time ranges (cluster-based permutation test, cluster-defining threshold p < 0.05, corrected significance level p < 0.05) for the 1^st^ (blue) and 2^nd^ (red) orientations.

### PING stimulus elicits backward reactivations during retention (Experiment 1)

After confirming the decoding approach during the encoding period, we next used the same analysis to examine the orientation representations held in WM that would be presumably reactivated by the PING stimulus during retention (see Figure 1A). Figure 3A plots the decoding performances for the 1^st^ (blue) and 2^nd^ (red) WM orientation features, as a function of time after the PING stimulus (see the reconstructed channel response in supplementary figure 3A). Interestingly, instead of being activated simultaneously, the 1^st^ and 2^nd^ orientations showed distinct temporal profiles, i.e., the 2^nd^ orientation showed earlier activation (T2: from 0.26 to 0.43 s, corrected cluster p = 0.011) than the 1^st^ orientation (T1: from 0.67 to 0.76 s, corrected cluster p = 0.030), with approximately 0.4 s temporal lag in-between. To further verify their distinct patterns, we computed the difference (Figure 3B) and sum (Figure 3C) between the 1^st^ and 2^nd^ decoding temporal profiles. The difference course was significant (0.31 – 0.42 s, corrected cluster p = 0.023; 0.62 – 0.75 s, corrected cluster p = 0.013) (Figure 3B), while their sum did not show any significance (Figure 3C), further confirming that the two items were activated at different latencies. The results suggest that the 1^st^ and 2^nd^ items, instead of residing in equally excitable states, tend to be stored in different latent states of the WM network. As a consequence, a transient perturbation of the network would produce an early 2^nd^ item reactivation and a late 1^st^ item response.

**Figure 3:**
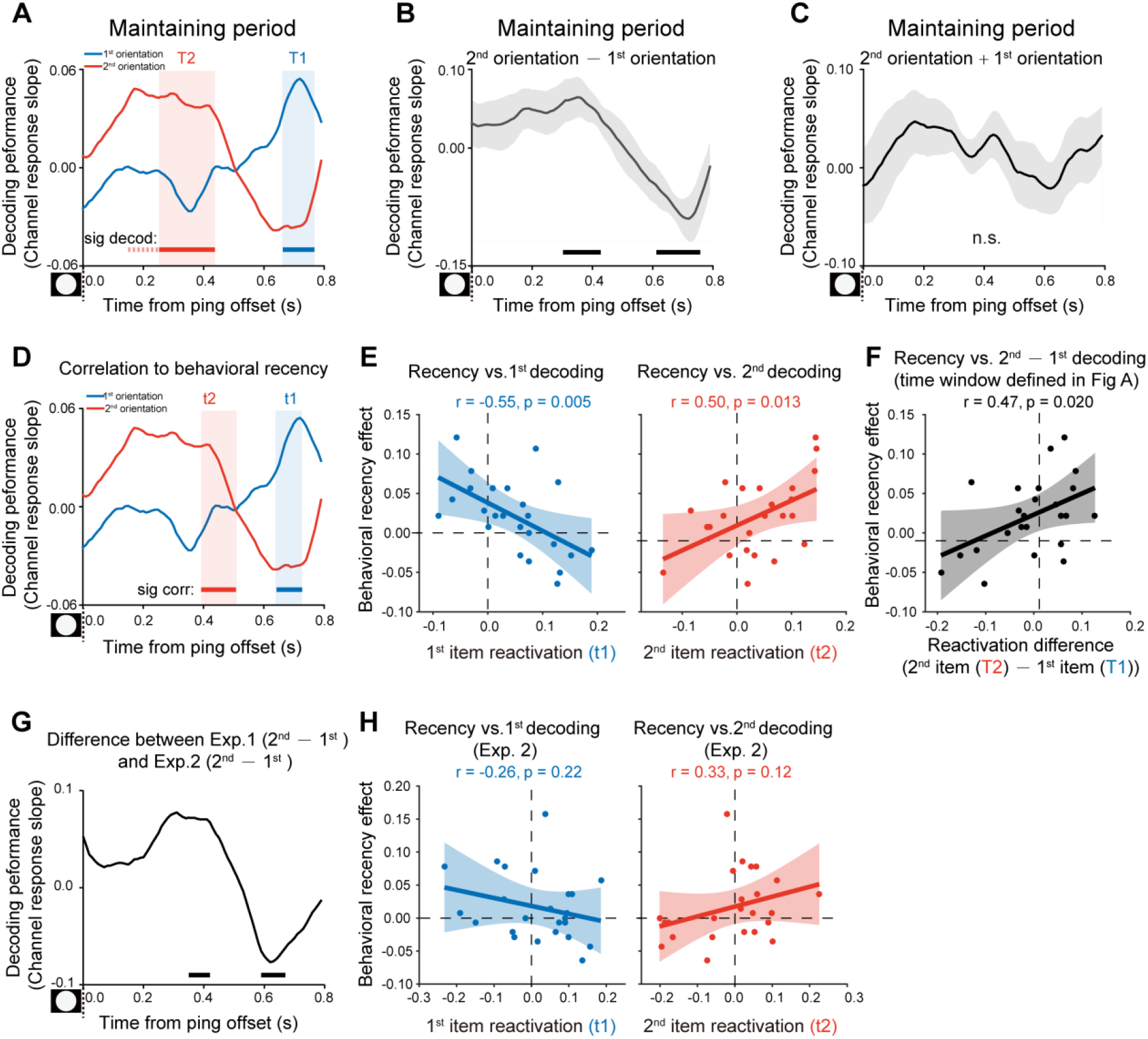
Backward reactivations during retention and behavioral relevance (Experiment 1). (A) Grand average time courses of the channel response slope for the 1^st^ (blue) and 2^nd^ (red) WM orientations after PING (inset in the bottom left) during the delay period. Horizontal lines below indicate time ranges of significant decoding strengths for the 1^st^ (blue, T1) and 2^nd^ (red, T2) orientations, respectively. (B) Grand average (mean ± SEM) time course of the difference between the 1^st^ and 2^nd^ channel response slopes (2^nd^ – 1^st^) and the significant time points. (C) Grand average (mean ± SEM) time course of the sum of the 1^st^ and 2^nd^ channel response slopes (1^st^ + 2^nd^). (D) Subject-by-subject correlations between the decoding performance and the recency effect were calculated at each time point. Horizontal lines below indicate time points of significant behavioral correlations (Pearson’s correlation after multi-comparison correction) for the 1^st^ (blue) and 2^nd^ (red) items, respectively. (E) Left: scatterplot (N = 24) of recency effect vs. 1^st^ decoding strength averaged over t1 (0.67 – 0.72 s after PING). Right: scatterplot of recency effect vs. 2^nd^ decoding strength averaged over t2 (0.4 – 0.43 s after PING). (F) Scatterplot (N = 24) of recency effect vs. decoding difference (2^nd^ at T2 – 1^st^ at T1). Each dot represents an individual subject. (G) Grand average time course of the 2^nd^ – 1^st^ reactivation difference between Exp 1 and Exp 2. (H) Same as E but for Exp. 2. (Horizontal solid line: cluster-based permutation test, cluster-defining threshold p < 0.05, corrected significance level p < 0.05; Horizontal dashed line: marginal significance, cluster-defining threshold p < 0.1, 0.05 < cluster p < 0.1. Shadow indicates 95% confidence interval).

### Reactivation profiles correlate with recency effect (Experiment 1)

We next evaluated the behavioral relevance of the reactivation profiles on a subject-by-subject basis, by relating the decoding strength to the recency effect, at each time point. As shown in Figure 3D, both the 1^st^ (blue) and 2^nd^ (red) item reactivations significantly correlated with the recency effect (horizontal lines, blue for the 1^st^ item and red for the 2^nd^ item), but at different time (1^st^ item: 0.65 – 0.72 s, corrected cluster p = 0.055, blue shades; 2^nd^ item: 0.4 – 0.5 s, corrected cluster p = 0.038, red shades) (see the raw correlation coefficient time course in supplementary figure 4A). Note that the temporal windows (t1, t2 in Figure 3D) showing significant neural-behavioral correlations was defined independent of the reactivation strength (Figure 3A). Figure 3E illustrates the correlation scatterplots within the two time windows (i.e., t1, t2) that were defined in Figure 3D, respectively. Specifically, the recency effect covaried positively with the 2^nd^ item (Figure 3E, right panel; Pearson’s correlation, N = 24, r = 0.50, p = 0.013) and negatively with the 1^st^ item (Figure 3E, left panel; Pearson’s correlation, N = 24, r = -0.55, p = 0.005). Moreover, we chose time bins solely based on significant reactivations regardless of its relevance to recency effect (Figure 3A; T1 for the 1^st^ item, blue shades; T2 for the 2^nd^ item, red shades). As shown in Figure 3F, consistently, the 2^nd^ – 1^st^ reactivation difference was correlated with the recency effect (Pearson’s correlation, N = 24, r = 0.47, p = 0.02). Taken together, the backward reactivation profiles that index distinct latent states in WM, show strong relevance to memory behavior, i.e., stronger recency effect is accompanied by larger, early 2^nd^ item reactivation and weaker, late 1^st^ item reactivation during the delay period.

**Figure 4:**
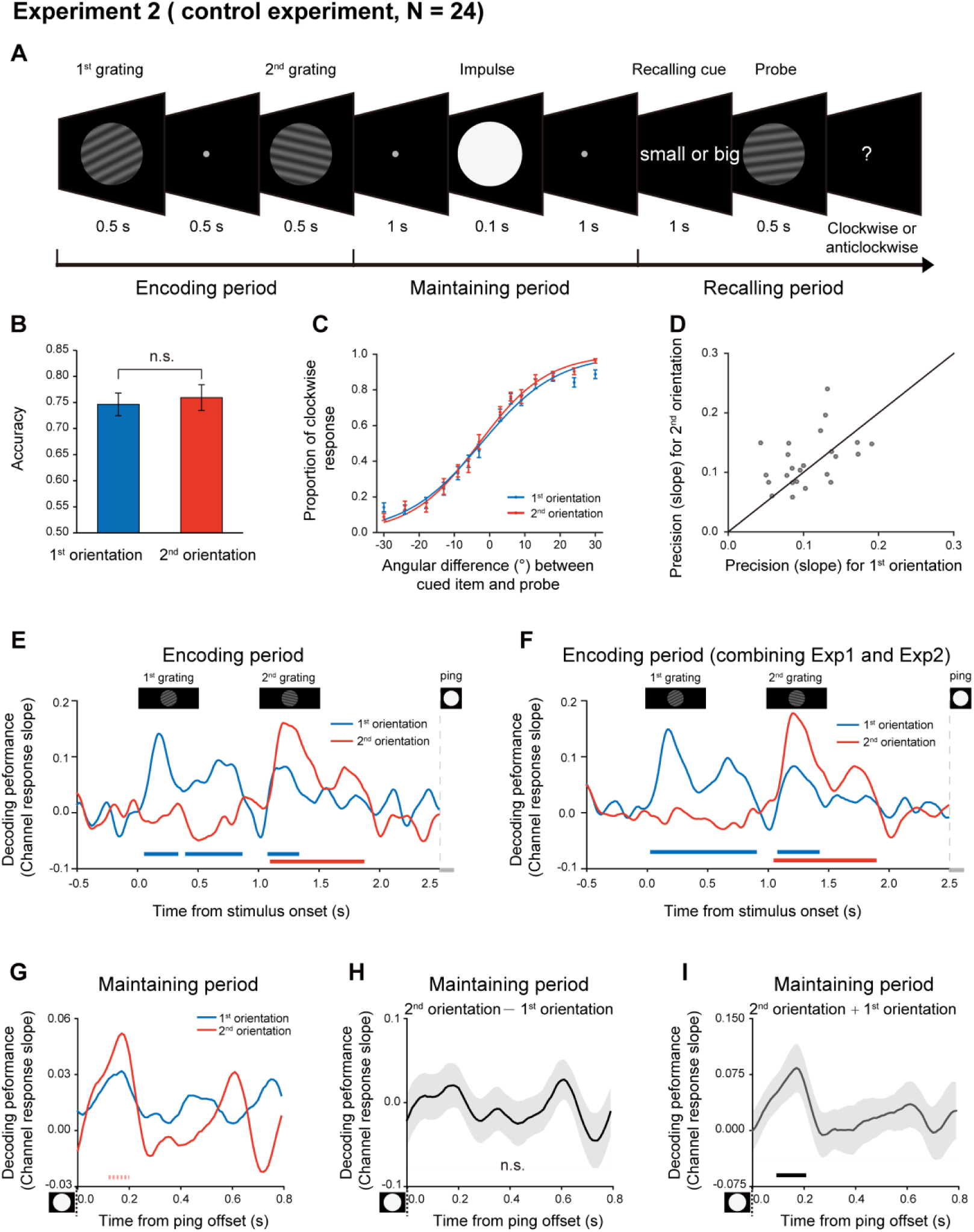
Experimental paradigm and results (Experiment 2). (A) Experiment 2 (N = 24) had the same stimuli and paradigm as Experiment 1, except that subjects did not need to retain the temporal order of the two orientation features. Subjects were first sequentially presented with two grating stimuli (1^st^ and 2^nd^ gratings) and needed to memorize the two orientations. During the recalling period, a retro-cue (small or big in character) appeared to instruct subjects which item that has either smaller or larger angular values relative to the vertical axis in a clockwise direction would be tested later. Next, a probe grating was presented and participants indicated whether the orientation of the probe was rotated clockwise or anticlockwise relative to that of the cued grating. Grand average (mean ± SEM) behavioral accuracy for the 1^st^ (blue) and 2^nd^ (red) orientations. Grand average (mean ± SEM) psychometric functions for the 1^st^ (blue) and 2^nd^ (red) orientations as a function of the angular difference between the probe and cued orientation. (D) Scatter plot for the slope of the psychometric function for the 1^st^ (x axis) and 2^nd^ (y axis) orientations. (E) Grand average time courses of the channel response slopes for the 1^st^ (blue) and 2^nd^ (red) orientations during the encoding period. Horizontal lines below indicate significant time points for the 1^st^ (blue) and 2^nd^ (red) orientations. (F) The same as E, but pooling Experiment 1 and Experiment 2 together during the encoding period (N = 48). (G) Grand average time courses of the channel response slope for the 1^st^ (blue) and 2^nd^ (red) WM orientations after PING (inset in the bottom left) during retention. (H) Grand average (mean ± SEM) time course of the difference between 1^st^ and 2^nd^ channel response slopes (2^nd^ – 1^st^). (I) Grand average (mean ± SEM) time course of the sum of the 1^st^ and 2^nd^ channel response slopes (1^st^ + 2^nd^) and significant time points. (Horizontal solid line: cluster-based permutation test, cluster-defining threshold p < 0.05, corrected significance level p < 0.05; Horizontal dashed line: marginal significance, cluster-defining threshold p < 0.1, 0.05 < cluster p < 0.1).

### Experimental procedure and disrupted recency effect (Experiment 2)

One possible reason for the backward reactivation in Experiment 1 is that the 2^nd^ item enters the memory system later than the 1^st^ item and thus decays more slowly, leading to a more excitable state for the lately presented item. In other words, the distinct latent states might solely arise from their different passive memory traces left in the network. To address the issue, we performed Experiment 2 using the same stimuli and paradigm as Experiment 1, except that subjects did not need to memorize the temporal order of the two orientations. Specifically, as shown in Figure 4A, subjects viewed two serially presented gratings and were instructed to memorize the two orientations.

During the recalling period, instead of indicating the 1^st^ or 2^nd^ orientation subjects need to recall as in Experiment 1, a retro-cue appeared to instruct subjects which item that has either smaller or larger angular values (relative to a vertical axis in a clockwise direction) would be tested later. Next, a probe grating was presented and participants indicated whether the orientation of the probe was rotated clockwise or anticlockwise relative to the cued grating. Thus, if the different latent states are due to the passive decay of the serially presented items, we would expect similar recency effect as well as backward reactivation as shown in Experiment 1.

Twenty-four new subjects participated in Experiment 2. Interestingly, the 1^st^ and 2^nd^ items showed similar memory performance (Figure 4B; 1^st^ item: mean = 0.77, s.e. = 0.011; 2^nd^ item: mean = 0.78, s.e = 0.011; paired t-test, df = 23, t = 1.57, p = 0.13, Cohen’s d = 0.32), and the psychometric functions did not differ in slopes for the 2^nd^ and 1^st^ orientations (Figure 4CD; N = 24, 1^st^ item: mean = 0.11, s.e. = 0.008; 2^nd^ item: mean = 0.12, s.e = 0.009; paired t-test, df = 23, t = 1.38, p = 0.18, Cohen’s d = 0.28). Thus, recency effect tends to be disrupted when the ordinal structure is not needed to be retained in WM.

### Disrupted backward reactivations during retention (Experiment 2)

We next used the same IEM approach to reconstruct the time-resolved neural representations of orientation features at each time point in Experiment 2. First, the encoding period showed a similar pattern as Experiment 1 (Figure 4E; also see the reconstructed channel response in supplementary figure 3BC). Specifically, the decoding performance of the 1^st^ (blue) and 2^nd^ (red) orientations displayed successive temporal profiles that were time-locked to the presentation of the corresponding grating stimuli (1^st^ item: 0.07 – 0.32 s, corrected cluster p = 0.005; 0.41 – 0.85 s, corrected cluster p = 0.002; 1.09 – 1.32 s, corrected cluster p = 0.005; 2^nd^ item: 1.11– 1.86 s, corrected cluster p < 0.001). Combining Experiment 1 and Experiment 2 revealed a clear serial profile during the encoding period (Figure 4F; 1^st^ item: 0.04 – 0.89 s, corrected cluster p < 0.001; 1.09 – 1.41 s, corrected cluster p = 0.003; 2^nd^ item: 1.06 – 1.88 s, corrected cluster p < 0.001).

In contrast, the reactivation profiles during the delay period in Experiment 2 did not show the backward pattern as observed in Experiment 1. Instead, the decoding performance for the 1^st^ and 2^nd^ WM orientations displayed similar temporal courses (Figure 4G; 0.1 – 0.2 s; 1^st^ item, p = 0.056, one-tailed; 2^nd^ item, p = 0.019, one-tailed). Consistently, the difference course between the 1^st^ and 2^nd^ decoding performances did not reach any statistical significance (Figure 4H; corrected cluster p > 0.5). Meanwhile, different from Experiment 1 (Figure 3C), the sum of the 1^st^ and 2^nd^ decoding performance showed significant responses (Figure 4I; 0.1 – 0.2 s, corrected cluster p = 0.024), further supporting their common reactivation profiles. As shown in Figure 3G, the 2^nd^ – 1^st^ reactivation difference (i.e., backward reactivation index) differed between Experiment 1 and 2, within the two temporal windows (0.35 – 0.42 s, independent t-test, cluster p = 0.058; 0.59 – 0.67 s, independent t-test, cluster p = 0.043). Furthermore, Experiment 2 did not show the reactivation-recency correlations either (Figure 3H), compared to Experiment 1 (supplementary figure 4B).

Since Experiment 2 instructed subjects to maintain the two orientations in terms of ‘big’ or ‘small’ labels, WM representations might be organized based on different principles (i.e., big or small) rather than ordinal position in Experiment 1. However, decoding analysis based on the big/small labels in Experiment 2 revealed similar reactivation profiles (supplementary figure 1), suggesting that the new labelling could not reorganize items in varied latent states as sequence structure context does. Finally, the neural representation of WM items could neither be generalized from the encoding to maintaining periods, nor across items in the reactivations triggered by PING stimulus (supplementary figure 2), further advocating that WM items embedded in the sequence structure context are reorganized in varied latent states of the WM system.

Taken together, when the two serially presented items are maintained in WM without a sequence structure imposed on them, they tend to be stored in comparable latent states of WM network, thereby having similar probability to be activated after a transient impulse and showing common reactivation profiles and no associations to the recency behavior. The results thus exclude the alternative interpretation that it is the different passive memory decay that gives rise to the different latent states as observed in Experiment 1.

Experiment 3 and computational model

Given the inherent time lag between the two sequentially presented items, the passive memory decay is seemingly a very straightforward interpretation for the recency effect. To further characterize the recency effect in terms of passive memory decay and sequence structure, we performed a behavioral experiment on 17 new subjects. Specifically, similar experiment design and task (retaining two orientations as well as their temporal order) as Experiment 1 (Figure 1A) were employed, except that there were 3 time intervals between the 2^nd^ grating (1 s, 2.5 s, 4 s) and PING stimulus. Notably, Experiment 1 with fixed interval between the 2^nd^ item and PING would make it difficult to reliably estimate the passive memory decay rate in behavioral performance, while the current design with three time lags would allow us to examine the passive memory decay and sequence context modulation in parallel from the same data set. As shown in Figure 5A, significant main effects in both serial position (i.e., recency effect; F_(1,16)_ = 10.41, p = 0.005, η^2^ = 0.39) and memory decay (F_(2,16)_ = 4.07, p = 0.027, η^2^ = 0.20) were observed in the behavioral experiment, and there was no interaction effect (F_(1,16)_ = 1.16, p = 0.33, η^2^ = 0.068). The results thus confirm that memory performance of items in a list would be determined by both passive memory decay and their positions in the sequence (e.g., 1^st^ or 2^nd^).

**Figure 5:**
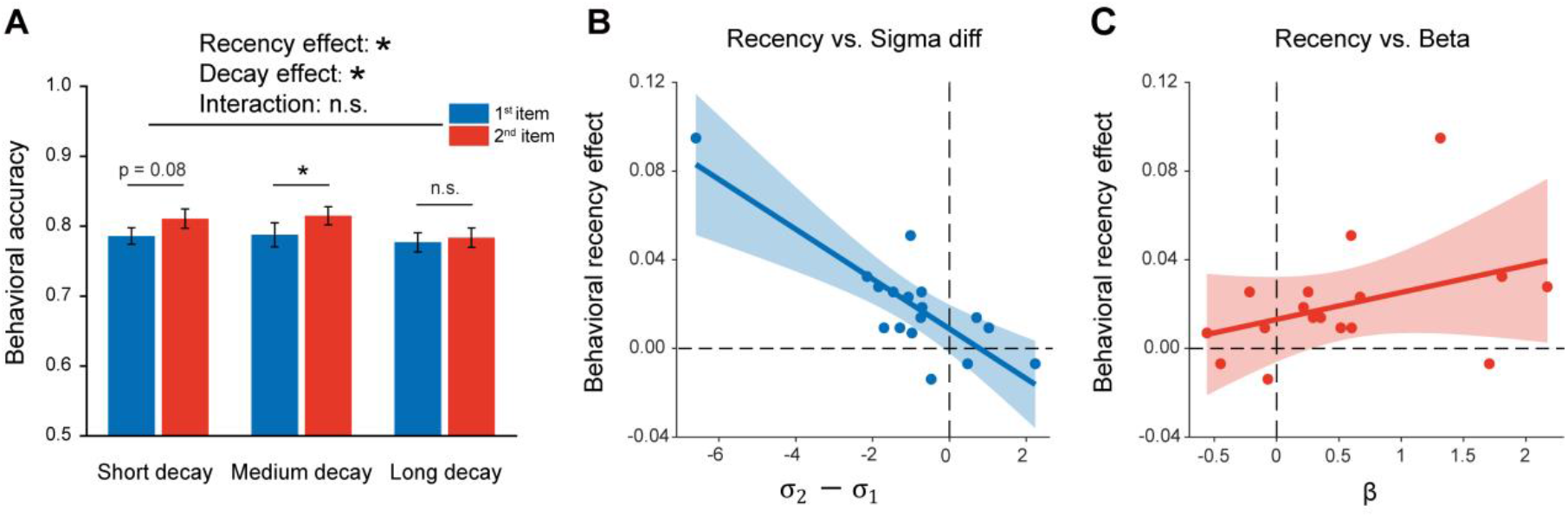
Experiment 3 and model fitting. Experiment 3 had the same paradigm as Experiment 1, but with 3 levels of 2^nd^ orientation-to-PING time intervals so that passive memory decay could be estimated. The model, endowed with sequence structure (*σ*_1_, *σ*_2_) and passive memory decay (β), was used to fit the behavioral data. (A) Behavioral results (N = 17). Grand averaged (mean ± SEM) behavioral accuracy for the 1^st^ (blue) and 2^nd^ (red) item at different 2^nd^ orientation-to-PING time intervals. (*: Two-way repeated ANOVA (Recency × Decay), p < 0.05). (B) Sequence structure (*σ*_2_− *σ*_1_) vs. recency effect (partial correlation, r = -0.82, p < 0.001). (C) Passive memory decay (β) vs. recency effect (partial correlation, r = 0.36, p = 0.17). The results support that the recency effect mainly derives from the sequence structure rather than passive memory decay.

We next built a computational model that comprises sequence structure (*σ*_1_and *σ*_2_for the 1^st^ and 2^nd^ items, separately) and passive memory decay (β) to assess their respective contribution to the recency effect. Specifically, for an item at a given time t after being encoded into WM, the standard deviation of its representational noise was set to be σ+β *t. The parameter β represents the memory decay rate and the parameter *σ*refers to the initial standard deviation of orientation representational noise at t = 0, whose value is either *σ*_1_(1^st^ item) or *σ*_2_(2^nd^ item). Since the 1^st^ item appears prior to the 2^nd^ item, it would have a longer *t* and in turn undergoes larger representational decay than the 2^nd^ item, presumably leading to the recency effect. On the other hand, *σ*_1_and *σ*_2_signify the abstract structure that organizes WM items by assigning different representational precision to items at different positions of a sequence (*σ*1for the 1^st^ item, *σ*2for the 2^nd^ item; lower value indicates higher representation precision). Thus, both the passive decay and sequence structure would presumably contribute to the recency effect, characterized by *β* and *σ*_2_− *σ*_1_, respectively.

The computational model was then fitted to the behavioral data to estimate the parameters (*β, σ*_1_, *σ*_2_) in each participant. A partial correlation analysis was then performed between the estimated parameters (*β,σ*_2_− *σ*_1_) and the recency effect across subjects. As shown in Figure 5, the sequence structure (*σ*_2_− *σ*_1_) was significantly correlated with recency effect (Figure 5B; r = -0.82, p < 0.001), while the decay rate (*β*) was not (Figure 5C; r = 0.36, p = 0.17), suggesting that the recency effect is mainly characterized by the sequence structure effect. Therefore, consistent with the MEG findings, the results support the central function of sequence structure, which represents the ordinal information of to-be-memorized items, in mediating the equence memory behavior, i.e., recency effect, independent of passive memory decay.

## Discussion

In two MEG experiments, we examined how the sequence structure imposed on a list of WM items shapes their neural representations in the human brain. Items located in different positions of a list are stored in distinct latent states of the WM neural network, being reactivated at different latencies. The reactivation pattern correlates with the recency effect in recognition behavior. In contrast, memorizing the same list of items without sequence structure requirements does not elicit the dissociated reactivations and displays nonsignificant recency effect. Moreover, neural representations of WM items could neither be generalized from the encoding to reactivations nor across items during retention, further advocating the reorganization of items in WM network. Finally, a computational model on the behavioral data supports that the recency effect is mainly derived from abstract sequence structures rather than passive memory decay. Taken together, sequence information, as a form of abstract structure context, essentially modulates memory performance by reorganizing items into different latent states of the WM neural network.

It has long been posited that WM relies on sustained neuronal firing or activities (Curtis & D’Esposito, 2003; Goldman-Rakic, 1995; Vogel & Machizawa, 2004). Interestingly, recent neural recordings and computational modeling advocate that memory could also be maintained in synapse weights without necessarily relying on sustained activities (i.e., hidden state), through short-term neural plasticity (STP) principles (Miller et al., 2018; Mongillo et al., 2008; Rose et al., 2016; Sprague et al., 2016; Stokes, 2015; Trübutschek et al., 2017; Wolff et al., 2017). Recent studies, by developing an interesting impulse-response approach, show that the neural representation of task-relevant features could be successfully reactivated from an activity-silent network (Wolff et al., 2017, 2020). Importantly, different levels of task relevance seem to be associated with different latent states these items are stored in, i.e., displaying reactivations at varied latencies. For instance, the immediately task-relevant item is in a more excitable state and tends to be first reactivated, whereas the item that is potentially task-relevant in the future resides in an activity-silent state and emerges later (e.g., Lewis-Peacock et al., 2011; Rose et al., 2016). Based on these findings, here we examined whether items in a sequence would be retained in varied latent state by examining their reactivation temporal profiles, i.e., one item stays in a more excitable state and tends to respond first. Importantly, the items in the current experiment are equally task-relevant and have the same probability to be tested during recalling. Thus, our findings could not be explained in terms of top-down, task-related attentional modulation as shown before.

In addition to top-down attentional modulation, low-level features could also modulate the excitability of neural populations. A theoretical model posits that multiple items, located in different excitable states according to their bottom-up saliency levels, would be reactivated at different phases within an alpha-band cycle (Jensen et al., 2012, 2014). Here, given the serial presentation of items during the encoding period, the lately presented item would presumably have less memory decay and in turn higher saliency level, hence residing in a more excitable state. However, two aspects of the results excluded the explanation. First, Experiment 2 used the same sequence of items as Experiment 1, yet did not reveal the temporally dissociated reactivation profiles. Indeed, when the sequence structure is not needed to be retained in WM, the two items are reactivated at similar latencies, indicating that they are stored in comparable latent states to be excited. Second, consistent with the idea, the computational modeling demonstrates that the passive memory decay could not reliably capture the recency effect; rather, it is the sequence structure that plays a central function in modulating recency effect.

An alternative interpretation for the backward reactivation is neural oscillations that are posited to mediate multi-item working memory (Bahramisharif et al., 2018; Huang et al., 2018; Lisman & Idiart, 1995; Lisman & Jensen, 2013; Roux & Uhlhaas, 2014). For example, previous studies demonstrate that items in a sequence fire spikes at different phase of neuronal oscillations (Siegel et al., 2009). Here, the time interval between the 1^st^ and 2^nd^ item reactivations is around 300-500 ms (Figure 3A), corresponding to the period of delta- and theta-band oscillations, thus suggesting that the observed sequential reactivation might derive from the low-frequency neuronal oscillations. Meanwhile, we only observed one round of sequential reactivation and thus could not make a strong case here. The lack of oscillatory reactivation profiles might be due to the gradually decreasing phase-locked neural reactivations following the PING stimulus, which would be even more difficult to be detected by noninvasive macroscopic-level MEG and EEG recordings.

Structure information has long been viewed to influence perception and memory, e.g., global precedence effect (Chen, 1982; L. Liu et al., 2017; Navon, 1977), reverse hierarchy theory (Ahissar & Hochstein, 2004), etc. Recently, it has been suggested that structural and content can be encoded in an independent manner to facilitate memory generalization (Behrens et al., 2018; Bengio et al., 2013; Higgins et al., 2017). Memory performance also tends to be influenced by the contexts the items are embedded in (DuBrow & Davachi, 2013; Gershman et al., 2013), i.e., when two items belong to different contexts, their temporal order relationship in memory would be largely disrupted. Objects that share the same contexts have been found to show stronger serial dependence across trials (Fischer et al., 2020). Furthermore, in a pure-tone frequency auditory discrimination task, subjects show a tendency to hold a perceptual anchor throughout the experiment (Harris, 1952; Mathias et al., 2020). In other words, the structure that characterizes the relationship between WM items spontaneously reshapes the representation of each item in WM. Meanwhile, not all types of structure could reorganize WM items, e.g., the decoding analysis based on big and small labels in Experiment 2 did not show the different reactivation profiles, suggesting that sequence structure serves as a special one. Thus, our results provide new converging behavioral, neural, and modeling evidence advocating the prominent influence of sequence structure on WM behavior and the underlying neural implementations – reorganization of items into different latent states.

The backward reactivation profile suggests that items located in the late position of a sequence are stored in a more excitable state, compared to those in the early position. The results are well consistent with our previous findings, whereby a temporal response function (TRF) approach was employed to tag the item-specific neural reactivations during retention in a sequence memory task. In that study, we observed backward reactivations, i.e., shorter latency for recent versus early items, which is further related to the recency effect (Huang et al., 2018). Meanwhile, the current findings differ from previous results in important ways. Specifically, the TRF latency refers to a relative time basis, whereas the present design assessed neural representations in absolute time after the PING, thus constituting new evidence for the backward reactivation. Moreover, the modeling as well as Experiment 2 further exclude the passive memory decay accounts for recency effect and backward reactivations, an unresolved issue in previous study (Huang et al., 2018). In fact, backward reactivations have been observed in many circumstances, e.g., during break after spatial experience in rats (D. J. Foster & Wilson, 2006), performing a reasoning task in humans (Kurth-Nelson et al., 2016), etc. A recent study shows that during structure learning, a forward replay in spontaneous activities would reverse in direction when paired with reward (Y. Liu et al., 2019), implying the involvement of reinforcement learning principles (Schultz et al., 1997; Sutton & Barto, 1998). Thus, a possible interpretation for the backward reactivation is that the item located in the late position of a list might serve as an anchoring point in memory for other items and is in turn maintained in a more excitable latent state, given that the recent item receives less inference and carries more reliable information for memory retrieval.

We built a computational model (Bays et al., 2009) that incorporates passive memory decay and sequence structure aiming to understand the recency behavior. Although both of the two accounts could in principle contribute to the recency effect, the model fitting on Experiment 3 separates the memory decay influence and confirms that it is the sequence context that mostly accounts for the recency effect. This is also consistent with a previous study revealing that low-level, absolute judgments fail to account for high-level, relative judgements (Ding et al., 2017). Although the current model only characterized the behavioral performance, the results are highly consistent with the MEG findings, i.e., decreased recency effect and comparable reactivations when sequence structure context was absent (Experiment 2). Thus, the MEG recordings together with the computational model constitute convergingly advocate a central function of sequence context in WM and its neural implementation in the human brain, i.e., reorganizing WM items into different latent states of the neural network.

## Author contributions

Q.H. and H.L. designed the experiment. Q.H. performed MEG and behavioral experiments and data analysis. H.Z. built the model. Q.H, H.Z., and H.L. wrote the paper.

## Acknowledgments

We thank Dr. Qing Yu and Dr. Ce Mo for helpful comments. We also thank the three reviewers for their important comments and suggestions in previous submission and the National Center for Protein Sciences at Peking University for assistance with MEG experiments. This work was supported by the National Natural Science Foundation of China (31930052 to H.L.), and Beijing Municipal Science & Technology Commission (Z181100001518002 to H.L.). Dr. Huihui Zhang was supported by Peking University Boya Postdoctoral Fellowship.

## Materials and Methods

### Ethics statement

Written informed consent was provided by all participants. Experimental protocols were approved by the Research Ethics Committee of Peking University and adhered to the principles conveyed in the Declaration of Helsinki.

### Participants

Twenty-four subjects (15 males, 21 ± 1.8 years old) participated in Experiment 1, and another 24 subjects (15 males, 21 ± 1.7 years old) participated in Experiment 2. Eighteen subjects (8 males, 21 ± 2.6 years old) participated in Experiment 3, and one subject was removed due to poor status. All subjects had normal or corrected-to-normal vision, with no history of psychiatric or neurological disorders. All subjects provided informed consent prior to the start of the experiment.

### Stimuli and tasks

#### Experiment 1

Each trial consisted of three phases – encoding, maintaining, and recalling. During the encoding period, participants were presented with two 0.5 s gratings (6° × 6°) sequentially at the center (0.5 s interval between them), and were instructed to memorize the orientations of the two gratings as well as their order, i.e., the 1^st^ orientation, the 2^nd^ orientation. For each trial, the orientations of the 1^st^ and 2^nd^ gratings were independently drawn from a uniform distribution over 25°– 175° in steps of 25°, plus a small random angular jitter (± 1° – ± 3°), with a constraint that they should differ at least by 25°. During the maintaining period, after 1 sec, a high luminance disc (30 cd/m^2^) appeared at the center for 0.1s, followed by another 1 s interval. During the recalling period, a retrospective cue (‘1’ or ‘2’ character) was first presented for 1 s to instruct subjects either the 1^st^ or 2^nd^ orientation would be tested. A probe grating (6° × 6°; 20% in contrast, 1 cycle per degree in spatial frequency, 2 cd/ m^2^ in mean luminance) was then presented for 0.5 sec at the center and participants indicated whether the orientation of the probe was rotated clockwise or anticlockwise relative to that of the cued grating. The angular differences between a memory item and the corresponding memory probe were uniformly distributed across seven angle differences (± 3°, ± 6°, ± 9°, ± 13°, ± 18 °, ± 24°, ± 30°). During each trial, all participants were instructed to keep the number of eye blinks to be minimum. Participants should complete 280 trials in total (determined in a pilot EEG study that used the same number of trials and found successful decoding), which took approximately 40 min (including breaks). Specifically, each subject completed 5 blocks with each of which containing 56 trials. The grating was chosen from 7 orientations and each orientation occurred 8 times per block with random order.

#### Experiment 2

Experiment 2 had the same stimuli and paradigm as Experiment 1, except that subjects did not need to retain the temporal order of the two orientation features. Specifically, in each trial, subjects were first sequentially presented with two grating stimuli and needed to memorize the two orientations without needing to retain their temporal order as in Experiment 1. During the maintaining period, a high luminance disc that did not contain any orientation information was presented. During the recalling period, a retro-cue appeared to instruct subjects which item that has either smaller or larger angular values relative to the vertical axis in a clockwise direction (‘big’ or ‘small’ in character) would be tested later. Next, a probe grating was presented and participants indicated whether the orientation of the probe was rotated clockwise or anticlockwise relative to that of the cued grating.

#### Experiment 3 for modelling (without MEG recording)

This experiment employed the same paradigm as Experiment 1, except that there were 3 temporal intervals between the offset of the 2^nd^ grating and the PING stimulus (i.e., 1 s, 2.5 s, 4 s). The orientations of the gratings were drawn from a uniform distribution over 20°– 170° in 30° increments, plus a small random angular jitter (± 1° – ± 3°), and the angular differences between a memory item and the corresponding memory probe were uniformly distributed across seven angle differences (± 3°, ± 6°, ± 9°, ± 13°, ± 18 °, ± 24°). Similar to Experiment 1, subjects were instructed to memorize the orientations of the two presented gratings as well as their order, i.e., the 1^st^ orientation, the 2^nd^ orientation. During the maintaining period, a high luminance disc (30 cd/m^2^) appeared at the center for 0.1s, followed by another 1 s interval. During the recalling period, a retrospective cue (‘1’ or ‘2’ character) was first presented for 1 s to instruct subjects either the 1^st^ or 2^nd^ orientation would be tested. A probe grating (6° × 6°) was then presented for 0.5 sec at the center and participants indicated whether the orientation of the probe was rotated clockwise or anticlockwise relative to that of the cued grating. Participants completed 864 trials in total, in three blocks.

### MEG recordings and preprocessing

Participants completed the MEG experiments inside a sound-attenuated, dimly lit, and magnetically shielded room. Stimuli were displayed onto a rear-projection screen (placed at a viewing distance of 75 cm) with a spatial resolution of 800 × 600 pixels and a refresh rate of 60 Hz. Neuromagnetic data were acquired using a 306-sensor MEG system (204 planar gradiometers, 102 magnetometers, Elekta Neuromag system, Helsinki, Finland) at Peking University, Beijing, China. Head movements across sessions should be within 3 mm for data to be involved for further analysis. The spatiotemporal signal space separation (tSSS) was used to remove the external noise (Taulu & Simola, 2006). Furthermore, both horizontal and vertical electrooculograms (EOGs) were recorded. MEG data were recorded at 1000 Hz sampling frequency. The MEG data was preprocessed offline using FieldTrip software (Oostenveld et al., 2011). Specifically, the data was offline band-pass filtered between 2 and 30 Hz. Independent component analysis was then performed in each subject to remove eye-movement and artifact components, and the remaining components were then back-projected to channel space. All data was then downsampled to 100 Hz. Trials with large variances or contained excessive or irregular artifacts were manually removed. Since here we focused on the orientation representations in the MEG response, only posterior MEG channels, including parietal sensors (52 planar gradiometers, 26 magnetometers) and occipital sensors (48 planar gradiometers, 24 magnetometers), were used for further analysis.

### Data analysis

#### Behavioral performance analysis

In addition to overall behavioral accuracy estimation, to further assess the psychometric function for orientation memory performance, we quantified the response proportion as a function of the angular difference between WM orientation and probe orientation. This function was further fitted in each subject, by *y* = 1/(1+ *e*^(−*β*\*(*x*−*μ*))^), where *β* represents the slope and *μ* is the bias parameter. The estimated slope *β* could represent memory precision, with larger value corresponding to better memory performance.

#### Time-resolved orientation decoding

To assess the time-resolved orientation information from the MEG signals, we used an inverted encoding model to reconstruct the orientation of the grating stimulus from the neural activities at each time point. This method has been previously used on many features, such as color (Brouwer & Heeger, 2009), orientation (Brouwer & Heeger, 2011; Ester et al., 2015; Kok et al., 2017; Myers et al., 2015), and spatial location (Sprague et al., 2014, 2016; Sutterer et al., 2019). One assumption of the model is that the response in each sensor could be approximated as a linear sum of underlying neural populations encoding different values of the feature-of-interest (e.g., orientation) separately, and therefore, by grouping the contributions from many sensors, we could achieve an estimation of the underlying neural population responses.

We began by modelling the response of each MEG sensor as a linear sum of seven information channels. *B*_1_(m sensors × n trials) represents the observed response at each sensor for each trial in the training set. *C*_1_(k channels × n trials) represents the predicted responses of each of the k information channels (i.e., k = 7 here) that are determined by basis functions, for each trial. *W* (m sensors × k channels) represents the weight matrix that characterizes the linear mapping from ‘channel space’ to ‘sensor space’. Taken together, their relationship could be described by a general linear model *B*_1_= *WC*_1_.

Specifically, similar to previous studies (Brouwer & Heeger, 2011; Ester et al., 2015), the basis functions that would determine *C*_1_are designed to contain seven half-wave rectified sinusoids centered at different orientation values (25°, 50°, 75°, and so on) and raised to the 6^th^ power. The weight matrix *W* (m sensors × k channels) could thus be estimated via using least-squares regression 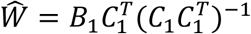.

After establishing *W* that links sensor space to the underlying information channels responses from the training set, we then used the estimated 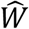 to test on independent datasets *B*_2_(sensors × trials) and calculated the predicted responses of the seven information channels, by 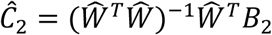.

The estimated channel responses Ĉ_2_was then circularly shifted to a common center (0°) in reference to the orientation-of-interest in each trial, which were further averaged across trials. A leave-one-out cross-validation was implemented such that data from all but one experimental block was used as *B*_1_to estimate 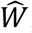, while data from the remaining block was used as *B*_2_to estimate *Ĉ*_2_, to ensure the independence between training set and testing set. The entire analysis was repeated on all combinations, and the resulting information channel responses were then averaged. Note that the procedure was performed at each time point so a time-resolved channel response for each subject was obtained.

To further characterize the orientation decoding performance, the slope of the calculated channel responses at each time was estimated by flipping the reconstruction performance across the center, averaging both sides, and performing linear regression (J. J. Foster et al., 2017). The slope time courses were further smoothed with a Gaussian kernel (s.d. = 40 ms, Wolff et al., 2017). The time-resolved decoding performance was tested by using one-sample t-test on the estimated slope against zero followed by multiple comparison correction across time (FieldTrip, cluster-based permutation test, 1000 permutations) (Maris & Oostenveld, 2007). All statistical tests were two-sided unless stated otherwise.

#### Correlations between recency behavior and neural reactivation profiles

To evaluate the behavioral relevance of neural reactivation profiles, we computed the Pearson’s correlation between the decoding performance and behavioral recency effect, at each time point, on a subject-by-subject basis. A multiple comparison correction across time was then performed on the correlation results by using cluster-based permutation test (N = 1000). To further illustrate the subject-by-subject correlations between the decoding strength and recency effect (Figure 3E), we averaged the decoding strengths over time-of-interests for the 1^st^ (t1: 0.67– 0.72 s, after PING) and 2^nd^ (t2: 0.4 – 0.43 s, after PING) items, respectively, in each subject.

The time-of-interests (i.e., t1, t2) were time points showing significant behavioral relevance as well as decoding strength. Note that the time-of-interests were just used for illustration purposes and the correlations between reactivations and the recency effect were independently tested with multiple-comparison correction over time, as described previously (Figure 3D).

We used another criterion to choose time-of-interests based on the decoding strength analysis (Figure 3A), i.e., T1 for the 1^st^ item (0.67 – 0.76 s, after PING) and T2 for the 2^nd^ item (0.26 – 0.43 s, after PING). The decoding strengths for the 1^st^ and 2^nd^ items were then averaged over T1 and T2, respectively, in each subject. The 2^nd^ – 1^st^ reactivation strength was then correlated with the recency effect (Pearson’s correlation; Figure 3F).

### Computational modeling

We built a computational model to account for the observed recency effect and the memory decay effect in Experiment 3. The model consists of two sets of parameters (β and σ), which characterize passive memory decay and the contextual influence of ordinal position, respectively (Figure 5). For an item, at a given time t after being encoded into working memory, the standard deviation of its representation noise was assumed to be σ+β * t (in three blocks, t = 3.1, 4.6, 6.1, respectively for item 1; t = 2.1, 3.6, 5.1 for item 2). The parameter β represents the memory decay rate and the parameter *σ*is the initial standard deviation of orientation representation noise at t = 0, whose value can be σ_1_or σ_2_ depending on the temporal order of items.

Because of the circular nature of orientation, for a given orientation, the probability distribution of its representation in working memory was assumed to be a von Mises distribution, which is a circular analogue of the familiar Gaussian distribution.

The general form of von Mises distribution is:

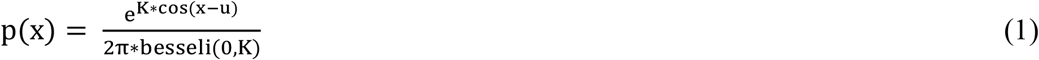

Here, x ranges from 0 to 2π. The parameter *u* is the mean of the distribution and the parameter K is a distribution shape parameter known as ‘concentration’. In our study, the orientation is a circular variable from 0 to π. In addition, we assumed that the representation precision did not differ for different orientations used in the current study. To simplify the calculation, all orientations of two items, *s*_1_and *s*_2_were set to 0. Thus, the probability distribution of their representation in working memory, p(x) can be written as:

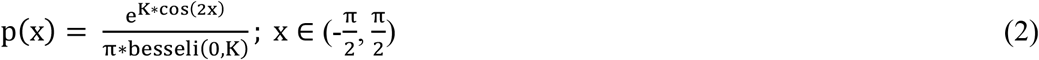

Conversion between the von Mises shape parameter K and the standard deviation of representation noise σ+β * t, K = sd2k(2 * (σ+β * t)) is achieved with sd2k function, which is adopted from Bays and his colleagues (Bays et al., 2009).

The probe orientation *s*_*p*_ is the relative orientation to the recalled orientation (*s*_1_or *s*_2_), ranging from 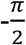 to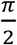 with positive values indicating clockwise compared with the recalled orientation. In each trial, for a given probe *s*_*p*_ (relative orientation), the probability of the binary choice (r = 1, correct; r = 0, wrong) is given by:

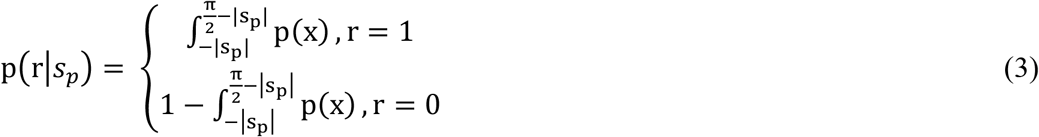

Assuming all trials are independent, the joint probability across all N trials can be written as:

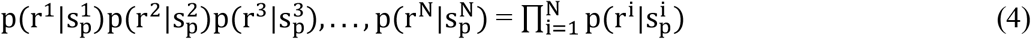

We next changed products to sums by taking the logarithm of both sides. The log likelihood of our model is given by:

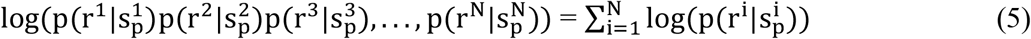

Together, we built a model of three parameters, β, σ_1_, σ_2_, to quantify the working memory of each item in a sequence. Here, β represents the decay rate of memory presentation precision. Parameters σ_1_and σ_2_reflect the contextual influence of ordinal position on the representation precision of items in a sequence.

The above model was then fitted to individual behavioral data. For each subject, parameters were estimated to produce the largest value of equation (5) using the Bayesian Adaptive Direct Search (BADS; Acerbi & Ma, 2017) with σ_1_, σ_2_∈ [1,50]; β ∈ [−5,5].

