## Supplementary for "Sequence structure organizes items in varied latent states of working memory neural network"

### Summmplementary figures

Supplementary figure 1

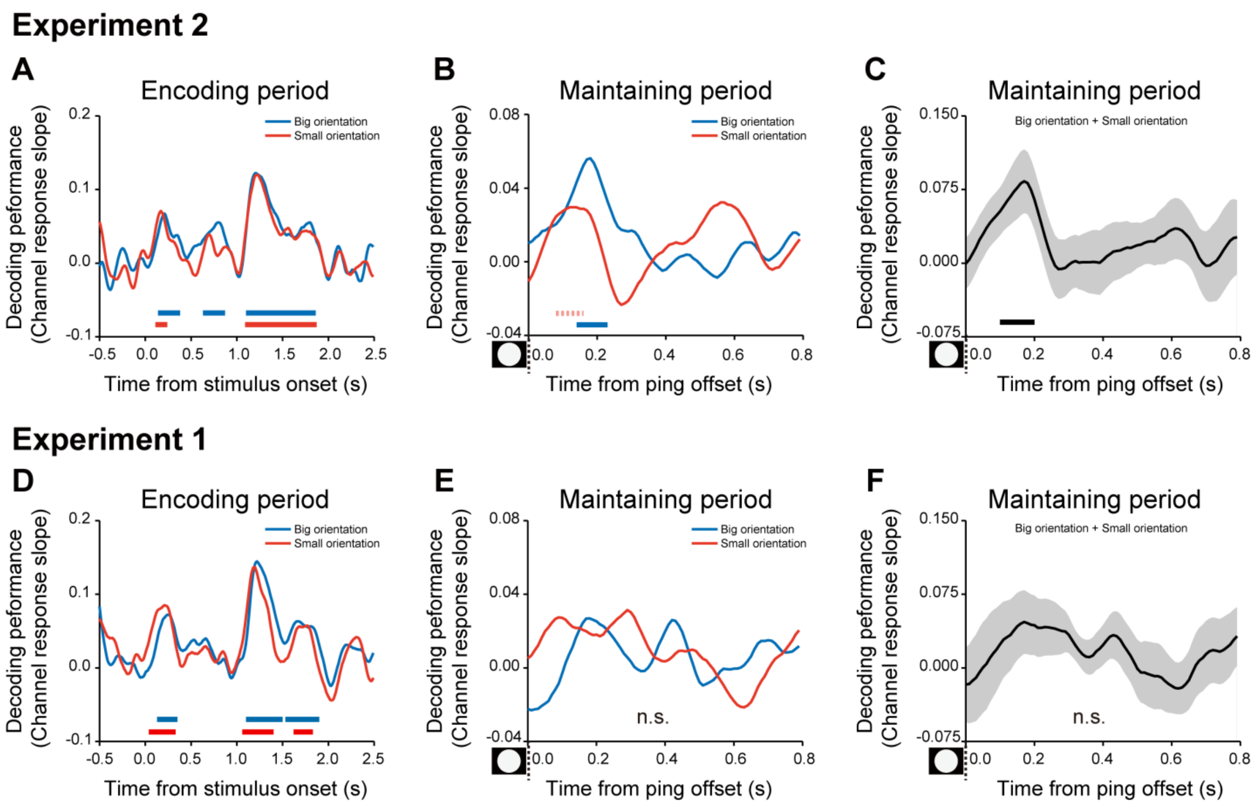


**Supplementary figure 1: Time-resolved orientation decoding performances based on big/small label for Exp.1 and Exp.2.**

(A-C) Exp.2. A: grand average time courses of the channel response slope for the 1$^{st}$ (blue) and 2$^{nd}$ (red) orientations during the encoding period. B: grand average time courses of the channel response slopes for the 1$^{st}$ (blue) and 2$^{nd}$ (red) orientations after the PING stimulus (inset in the bottom left) during the delay period. C: grand average (mean ± SEM) time course of the sum of the 1$^{st}$ and 2$^{nd}$ channel response slopes (1$^{st}$ + 2$^{nd}$) during the maintaining period. (D-F)The same as A-C, but for Exp.1. Horizontal solid line: cluster-based permutation test, cluster-defining threshold $p < 0.05$, corrected significance level $p < 0.05$; Horizontal dashed line: marginal significance, cluster-defining threshold $p < 0.1$, $0.05 <$ cluster $p < 0.1$.

Supplementary figure 2

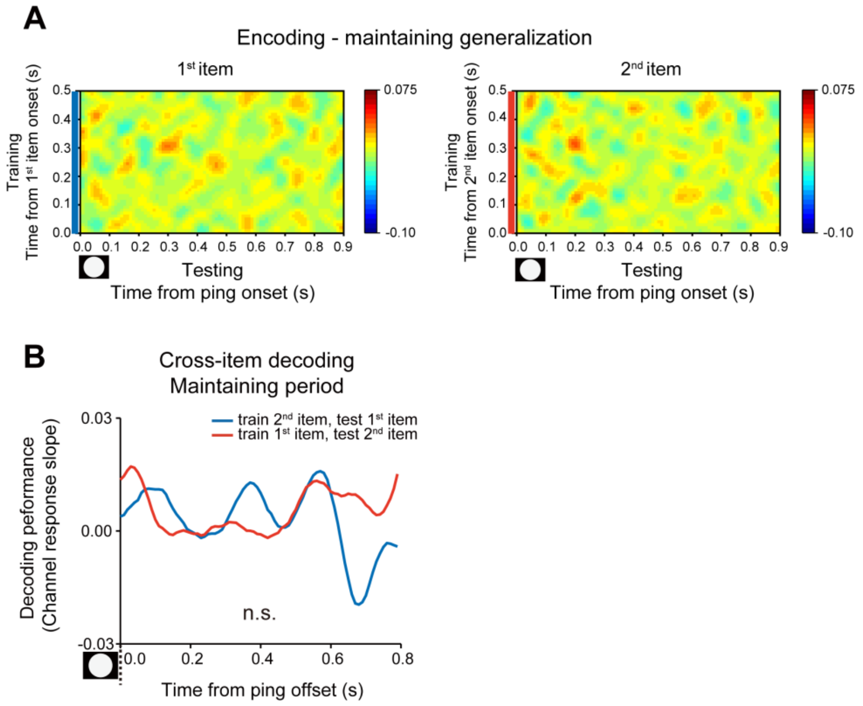


**Supplementary figure 2: Cross-generalization results (Exp 1)**

(A) Encoding-to-maintaining generalization matrices for the 1st (left) and 2nd (right) orientations by training decoders on each time point of the encoding period (y-axis) and testing all time points of the maintaining period (x-axis). (B) Cross-item generalization during the maintaining period. Blue line indicates training decoder based on the 2nd item and testing on the 1st item. Red line indicates training decoder based on the 1st item and testing on the 2nd item.

Supplementary figure 3

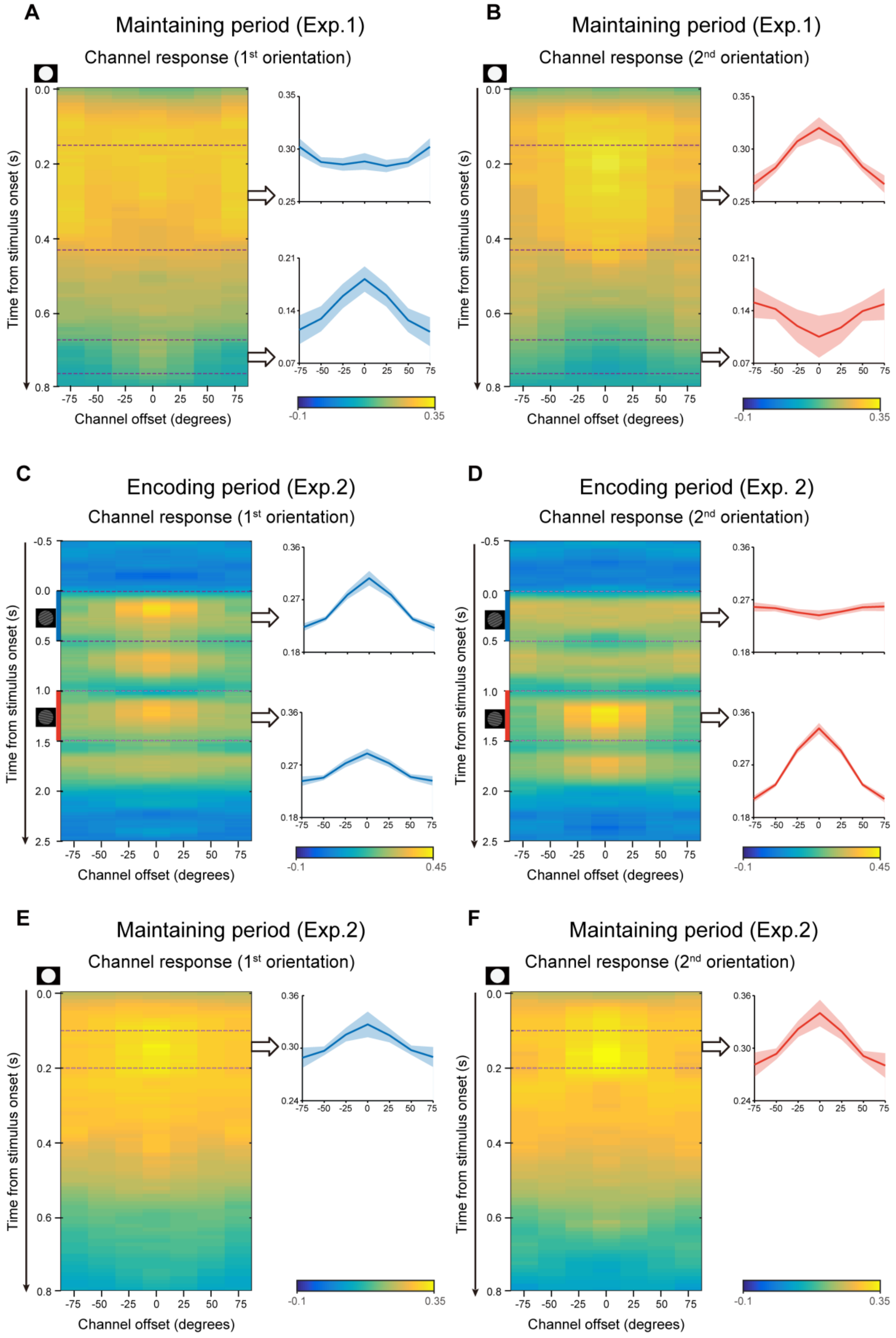

**Supplementary figure 3: Time-resolved reconstructed channel responses**. (A-B) Maintaining period in Exp.1. A: grand average time-resolved channel response for the 1st orientation after the PING stimulus during the delay period (left). Grand average (mean ± SEM) channel response for the 1st orientation averaged over the 0.15 – 0.43 (right upper) and 0.67 – 0.76 s (right lower). B: same as A, but for the 2nd item. (C-D) Encoding period in Exp.2. C: grand average time-resolved channel response for the 1st orientation throughout the encoding period. Grand average (mean ± SEM) channel response for the 1st orientation averaged over the 1st grating presentation period (0 – 0.5 s, right upper) and the 2nd grating presentation (1 – 1.5 s, right lower). D: same as C, but for the 2nd item. (E-F) Maintaining period in Exp.2. E: grand average time-resolved channel response for the 1st orientation during the maintaining period (left). Grand average (mean ± SEM) channel response for the 1st orientation averaged over the 0.1 – 0.2 s (right). F: same as E, but for the 2nd item.

Supplementary figure 4

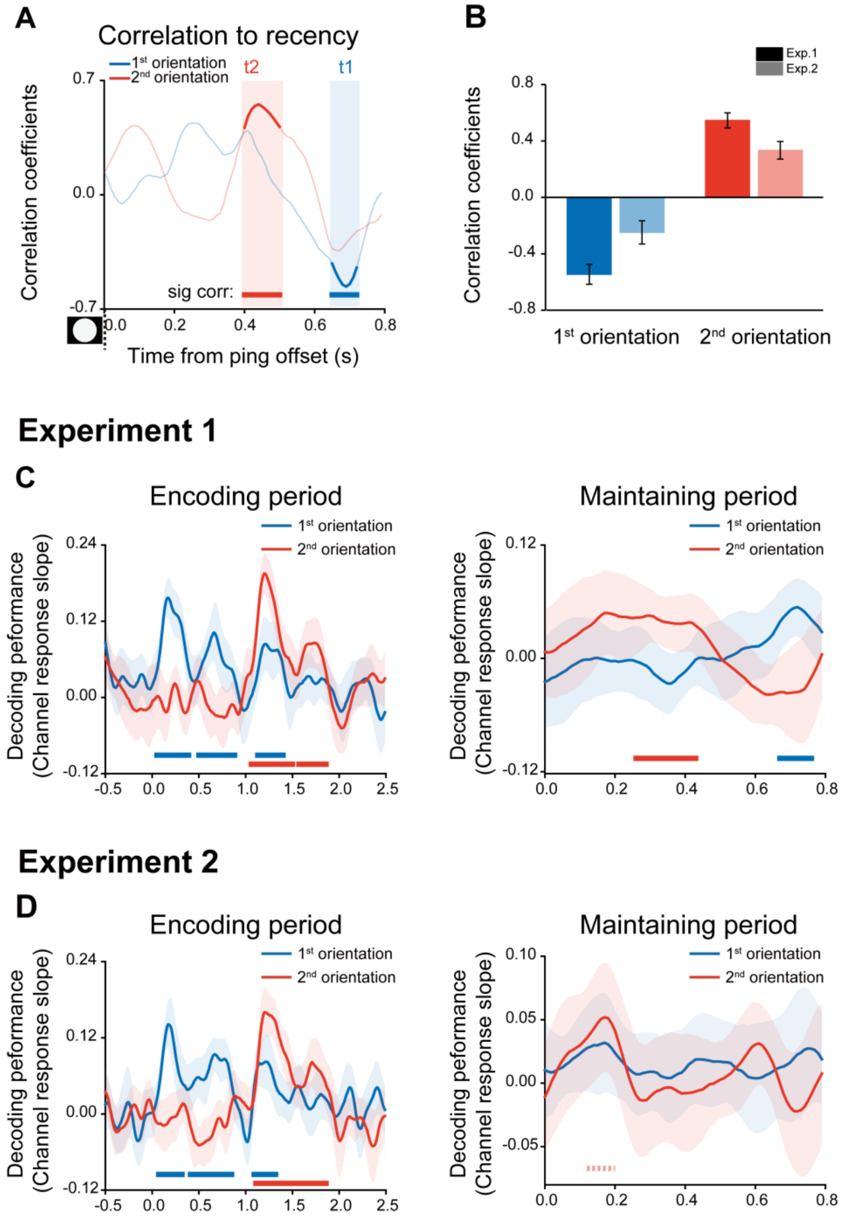


**Supplementary figure 4**

(A) Neural-recency correlation coefficients time courses in Exp.1. Horizontal lines below indicate time points of significant behavioral correlations (Pearson's correlation after multi-comparison correction) for the 1st (blue) and 2nd (red) items, respectively. (B) Correlation coefficients between recency and reactivation strength (Mean ± confidence interval) for 1st (t1, dark blue for Exp.1, light blue for Exp.2) and 2nd items (t2, dark red for Exp.1, light red for Exp.2). The confidence intervals were calculated using a jackknife procedure. (C) Grand average (Mean ± 95% confidence interval) time courses of channel response slope for the 1st (blue line) and 2nd (red line) orientations during encoding (left) and maintaining period (right) in Exp.1. (D) The same as C, but for Exp.2. (Horizontal solid line: cluster-based permutation test, cluster-defining threshold $p < 0.05$, corrected significance level $p < 0.05$; Horizontal dashed line: marginal significance, cluster-defining threshold $p < 0.1$, $0.05 <$ cluster $p < 0.1$).
